# NordicTraits: imputed species-level functional trait dataset for vascular plants of Denmark, Finland, Iceland, Norway and Sweden

**DOI:** 10.64898/2026.03.03.709463

**Authors:** Pekka Niittynen, Risto K. Heikkinen, Maria H. Hällfors, Aino-Maija Määttänen, Veera Norros, Julia Kemppinen

## Abstract

The NordicTraits dataset provides the first comprehensive, imputed, and openly available species-level functional trait resource for all native vascular plants across Denmark, Finland, Iceland, Norway, and Sweden. Functional traits such as plant height, seed mass, and leaf nitrogen content are critical for understanding plant strategies, ecosystem processes including the services they provide to human society, and predicting biodiversity responses to environmental change. The Nordic region has a rich botanical history. However, the absence of a unified trait database has limited trait-based ecological research in this region that is under rapid climate change. To address this gap, we compiled and harmonized trait data from major global databases and regional sources, covering 3,099 vascular plant species. We utilized all together 205 traits in the imputation model with the source data covering, on average, 54% (5–81%) of the species. We employed rigorous data cleaning, taxonomic standardization, and a Random Forest-based imputation framework to fill the missing values, while incorporating phylogenetic information to improve accuracy. The final dataset includes 44 selected key functional traits with no missing values, including both continuous and categorical traits and enabling robust analyses of plant strategies and responses to environmental gradients across the region’s diverse temperate, boreal, arctic, and alpine ecosystems. The dataset is particularly valuable for large-scale, multi-species studies, and those focusing on functional community assessments across a wide range of vegetation types. NordicTraits facilitates the paradigm shift from species-based to trait-based ecology, supporting research on biodiversity, conservation, and climate change impact predictions in northern Europe.

## 1. Introduction

Plant functional traits are morphological, physiological, phenological, or behavioral characteristics of plants that influence their performance, fitness, and role in ecosystems (Carlos M. Duarte et al. 1995, Diaz et al. 1998, 2016). These traits include features such as plant height, seed mass, leaf nitrogen content, and dispersal distance, which collectively reflect species’ resource acquisition strategies, competitive ability, and responses to environmental conditions and change (Kattge et al. 2020). Functional traits can be used to connect individual plant performance to ecosystem-level processes and provide a mechanistic understanding of how plants interact with their environment and with other organisms (Diaz et al. 2004, Violle et al. 2007, Webb et al. 2010).

Recently, the emergence of trait-based methods has flourished in plant ecology (Westoby 2025). Trait-based approaches can provide a powerful framework in ecology, enabling generalizations of ecological processes across different regions, scales, and ecological hierarchies, and ultimately allowing novel inferences beyond taxonomic classifications (Webb et al. 2010, Enquist et al. 2015). Furthermore, plant functional traits are instrumental in predicting biodiversity shifts from the individual to ecosystem level. Traits govern community assembly, influence invasion success, mediate the impact of species diversity on ecosystem responses and stability, and shape responses of communities to climate change (de Bello et al. 2010, 2021, Schob et al. 2013, Kraft et al. 2015, Funk et al. 2017, Westerband et al. 2021). Thus, understanding variation in functional traits across species allows scientists to better predict vegetation responses to different environmental changes. This highlights the critical role of trait data in ecological and evolutionary research, conservation planning, and ecosystem management (Hortal et al. 2015a, Carlucci et al. 2020, Miatta et al. 2021).

The Nordic region encompasses Denmark, Finland, Iceland, Norway, and Sweden, and is characterized by a high diversity of vegetation types (Lawesson and Skov 2002, Lindholm and Heikkilä 2006, Wasowicz et al. 2014, Virtanen et al. 2016). The Scandes mountain range forms a major geographic and climatic divide, separating the oceanic influences of the Atlantic from the continental climate of Eurasia (Pedersen 1990). This historical and ongoing climatic differentiation has profoundly shaped the region’s natural colonization history and its unique biodiversity, which comprises circumpolar, Atlantic, and Eurasian species (Kullman 2008, Seppä et al. 2009). Botanical research has a long tradition in the Nordic countries but less attention has been paid to trait-based ecology, and more importantly, no comprehensive trait database exists comparable to those compiled for the Mediterranean region (Tavşanoğlu and Pausas 2018), China (Wang et al. 2022) or Australia (Falster et al. 2021).

Here, our objective is to compile a comprehensive species-level plant functional trait dataset covering all native plant species across the diverse ecosystems of the Nordic countries and incorporating both continuous and categorical traits. This dataset is specifically designed to overcome the pervasive challenge of missing data in existing global trait databases through the application of advanced imputation methodologies. Ultimately, our dataset aims to serve as a taxonomically encompassing, robust, openly available, and accessible resource to support a wide array of ecological and evolutionary studies, facilitating a paradigm shift from traditional species-based ecology to a more mechanistic, trait-based understanding (Kattge et al. 2020). By employing multiple data sources, sophisticated data cleaning, and a robust Random Forest-based imputation, the NordicTraits dataset compiles fragmented, heterogeneous, and incomplete empirical data into a regionally complete and analytically powerful resource. The broad coverage of species functional traits allows developing an in-depth mechanistic understanding of plant strategies and their responses to environmental gradients and biotic pressures across the diverse Nordic ecosystems.

## 2. Materials and Methods

The development of the NordicTraits dataset followed a multi-stage workflow designed to make a leap from heterogeneous raw trait observations to a harmonized, gap-free species-level trait matrix. Our approach integrated taxonomic standardization for 3,099 vascular plant species occurring in the Nordic countries, systematic data mining across 20 trait databases, and a phylogenetically-informed imputation framework using Random Forests. This systematic progression, encompassing initial data filtering, species-level aggregation, iterative imputation, and rigorous quality validation, is summarized in **Figure 1**. All data processing was conducted in R 4.5.1 (R Core Team 2025).

**Figure 1.**
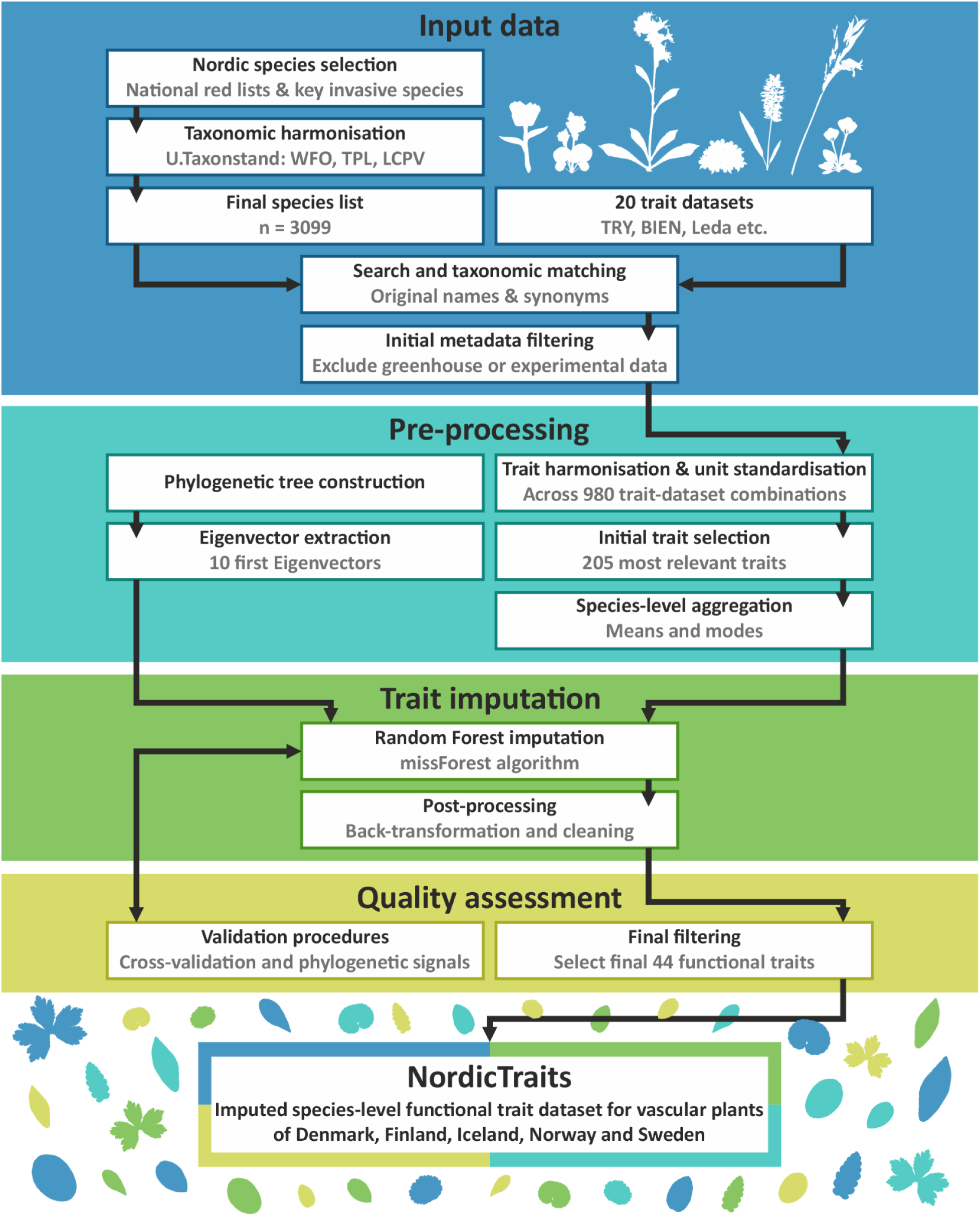
The workflow for generating the NordicTraits dataset.

### 2.1 Species list and taxonomic harmonization

We constructed a species list of vascular plants native to the five Nordic countries by combining the national assessments of threatened species (Hyvärinen et al. 2019, Wasowicz and Heiðmarsson 2019, SLU Artdatabanken 2020, Artsdatabanken 2021, Moeslund et al. 2023). The Norwegian assessment includes species in Svalbard but the Danish species list does not cover species of Greenland or Faroe Islands. We included all vascular plant species that were considered resident and have been assessed in any of the countries. Additionally, we included all species in the trait dataset by Tyler at al. (2021) which also includes the most important invasive species and some of the most commonly occurring non-resident species in Sweden. We consider that this list represents the key non-resident species also across other Nordic countries.

We used the *nameMatch* function in the *U.Taxonstand* R package (Zhang and Qian 2023, Zhang et al. 2025) to standardize taxon names across the country-level species lists with three global taxonomic databases, namely The Plant List (The Plant List 2013), The World Flora Online (WFO 2025), and The Leipzig Catalogue of Vascular Plants (Freiberg et al. 2020). We included the taxon author information when available to increase the accuracy of the name matching and synonym searching. We screened the name matching results manually and decided to use the World Flora Online (WFO) names as our taxonomic backbone in later steps as it had the least inconsistencies and found a resolved name most often. However, we manually checked, searched, and corrected all taxa for which the *nameMatch* function did not provide a matching name in WFO database or when the three databases provided diverging output. For example, the *nameMatch* function allows fuzzy matching (i.e., allows small differences in species names) and thus led to some errors when an exact match was not found. After the taxonomic standardization we arrived at a final list of 3099 Nordic plant species.

### 2.2 Compiling the trait data

The NordicTraits plant functional trait dataset was compiled using various major trait databases as input for trait observations (i.e., individual species-level trait measurements). The databases that were used included, e.g., TRY Plant Trait database (Kattge et al. 2020), Botanical Information and Ecology Network (BIEN; Enquist et al. 2016, Maitner et al. 2018), the LEDA Traitbase (Kleyer et al. 2008), FloraVeg.EU database (Chytrý et al. 2024), Tundra Trait Team database (TTT; Bjorkman et al. 2018), Fine-Root Ecology Database (FRED; Iversen et al. 2021), Global Inventory of Floras and Traits (GIFT; Weigelt et al. 2020), and Global Root Trait Database (GRoot; Guerrero-Ramirez et al. 2021). In addition, we searched for additional datasets and research articles with accompanying original trait data covering northern European flora using Google Dataset Search, and the Zenodo and Dryad data repositories (see a summary of all data sources in **Table 1**). We used the BIEN (Maitner et al. 2018), GIFT (Denelle et al. 2023), and TR8 (Bocci 2015) R packages to access the corresponding databases. The rest were downloaded and compiled manually. From hereon, we refer to the main data sources and databases as datasets and the original sources (e.g., an original scientific study) as subdataset – e.g., the full TRY database is a dataset and all the primary data providers within it are called subdatasets.

**Table 1.** Sources of trait observations. N_obs = total number of trait observations for the species on our list (3099 species); N_traits = number of traits included in the dataset; N_species = number of plant species in the dataset that correspond to our species list (3099 species).

| Dataset | N_obs | N_traits | N_species | Reference |
| --- | --- | --- | --- | --- |
| BIEN | 3093365 | 24 | 2313 | (Enquist et al. 2016, Maitner et al. 2018) |
| Baruah | 1498 | 3 | 28 | (Baruah et al. 2017) |
| Drevojan | 35256 | 13 | 2262 | (Dřevojan et al. 2023, Midolo et al. 2024) |
| FRED | 35914 | 16 | 916 | (Iversen et al. 2021) |
| GIFT | 275368 | 50 | 2563 | (Weigelt et al. 2020) |
| GRoot | 39437 | 15 | 1467 | (Guerrero-Ramirez et al. 2021) |
| LEDA | 105622 | 24 | 2054 | (Kleyer et al. 2008) |
| Lososova | 11020 | 5 | 2204 | (Lososová et al. 2023) |
| Majekova | 549 | 6 | 99 | (Májeková et al. 2021) |
| Midolo | 11304 | 6 | 1884 | (Midolo et al. 2023) |
| Mudrak | 11190 | 8 | 60 | (Mudrák et al. 2019) |
| Niittynen | 114350 | 5 | 356 | Data article in preparation, part of the data used in (Kemppinen and Niittynen 2022, Abrego et al. 2025) |
| TR8 | 53743 | 79 | 1895 | (Bocci 2015) |
| TRY | 1106167 | 124 | 2500 | (Kattge et al. 2011, 2020) |
| TTT | 57642 | 17 | 359 | (Bjorkman et al. 2018) |
| Tesitel | 2952 | 1 | 2952 | (Těšitel et al. 2024, Fahs et al. 2025) |
| Tichy | 13278 | 6 | 1990 | (Tichý et al. 2023) |
| Tyler | 141014 | 63 | 2275 | (Tyler et al. 2021) |
| Zanne | 2288 | 2 | 1704 | (Chave et al. 2009) |
| Zuijlen | 610 | 5 | 22 | (van Zuijlen et al. 2022) |

We used all the species names present in the original country-level species lists (see section 2.1) for the 3099 Nordic species to search for trait data in all the selected trait datasets. We conducted the searches including the species names with and without the author citation attached to the names. This extended species name list helped us maximize the number of trait observations for our species set as the used nomenclature and naming syntax varied greatly across the trait data sources. Next, we used the same taxonomic standardization process as for the country-level species lists described above to match all the species names across sources and to select the relevant trait data. Furthermore, we screened the metadata describing the trait observations and excluded all data collected from plants that had been grown in green houses or manipulated experimentally.

Here, we acknowledge that among the 3099 species, all together 600 species belonged to only three genera, namely *Taraxacum*, *Hieracium,* and *Ranunculus* which are known for their complex and often unresolved taxonomy. Due to this complex taxonomy, many of the trait data sources included information only at genus-level for these genera, and thus species-level trait observations were especially scarce within these three genera.

### 2.3 Preprocessing the trait data

We partitioned the compiled trait data based on their original source (i.e. subdataset), to facilitate a systematic data processing procedure for the purposes of constructing NordicTraits. This partitioning resulted in 980 combinations of individual traits and datasets. We manually inspected all the 980 combinations and identified which of these combinations effectively represented the same functional trait and harmonized them. For example, many original data sources had a trait describing the woodiness of the species but used different class names that needed to be harmonized before combining.

For each continuous trait, we transformed all observations to the same unit. We also screened for clear unit errors by calculating summary statistics for each subdataset and trait, enabling easy detection of subdatasets that were, e.g., systematically several magnitudes higher than values for the same species in other subdatasets, thus allowing correcting with high certainty. To further remove erroneous observations, we applied logical thresholds, i.e., theoretical minimum and maximum values for traits where such thresholds are applicable (e.g., by bounding leaf dry matter content between 0 and 1). When needed, we divided the continuous traits into separate traits based on how they were measured, for instance separating leaf area into two traits based on whether the petiole was included or excluded in the measurements. However, for many (sub-) datasets such detailed information on methodology was not available, and in such cases we classified the trait based on the most likely method used, thus potentially creating some noise in the data. We opted for making these assumptions and retaining the data, rather than removing such observations, since similar lack of methodological information was relatively frequent in the source data and would have resulted in discarding large amounts of observations if following a stringent inclusion approach.

For categorical traits, we harmonized the conceptually matching traits when possible (e.g., combined all traits describing woodiness of the species across sources) or divided them into separate traits based on the data source when harmonization of the used categories was not feasible. For instance, the same conceptual trait describing primary pollination mode sometimes followed a slightly different classification system in different source datasets. In such cases, we decided to retain both related traits rather than combining them to one trait with a broad definition, which would ultimately retain less information for the trait imputation step (see section 2.6). Additionally, some categorical traits were transformed into binary dummy variables to account for species possessing multiple attributes within a single trait (e.g., traits describing dispersal vectors).

Next, we made an initial selection of the traits based on two criteria: 1) traits that are most commonly used and referred to in trait-based ecological literature and 2) traits that had the highest coverage across our 3099 Nordic species. For the first criteria, we made exceptions for traits that – while not commonly included in trait-based assessments – describe ecologically important aspects of form or function for plants. For instance, compared to above-ground traits, root traits are generally less prevalent in the trait databases and scientific studies. Nevertheless, root traits are key for understanding many ecosystem functions, and with recent rising interests in understanding below-ground biodiversity and ecosystem functions, we considered these traits to be of high potential value. Albeit, as the NordicTraits database focuses on traits that are relevant across all plant species, we left out some group-specific traits such as wood-related traits. While our end-goal was to produce a dataset on traits related to form, function, and strategy, at this stage we also opted to include traits on biogeography and macroecology (such as Ellenberg values or habitat preferences) as such variables may provide important ecological context to the imputation model, especially for species with a large proportion of missing trait values.

These trait selection and harmonization steps resulted in 205 trait variables that were included in the imputation process. Of the 205 traits, root phosphorus content covered the lowest proportion of the species (5.1%) while photosynthetic pathway had the highest coverage (81.3%). On average, the traits covered 54% of the 3099 species (**Figure 2**).

**Figure 2.**
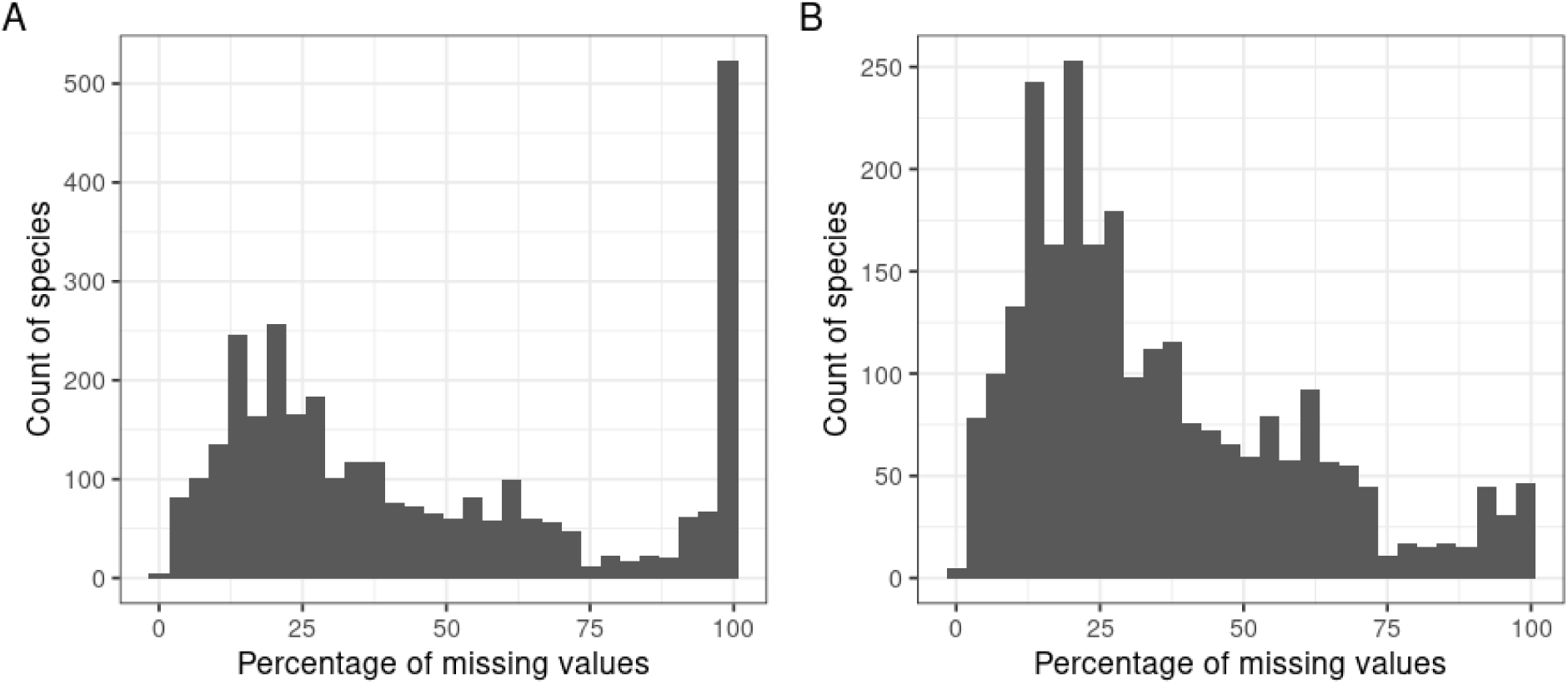
Distribution of percentage of missing values across the 205 initially selected traits for A) all the 3099 species, and B) for 2499 species excluding taxonomically complex genera Taraxacum, Hieracium and Ranunculus.

### 2.4 Aggregating species-level traits

In many cases, the trait datasets contained multiple measures per trait and per species, which represent intraspecific trait variation and measurement error. Here, our aim was to produce a comprehensive species-level trait dataset with no missing values, and therefore, we first aggregated the trait data to species-level values prior to imputation. This section describes the procedure for calculating the species-level trait values.

The number of trait observations varied greatly across (sub-)datasets (Table 1). Furthermore, some of the datasets provided aggregated species/population level averages while others provided individual-level measurements from specific studies. This means that the most observation-rich datasets would have dominated the calculation of species-level trait values if this issue would not have been considered, thereby introducing potential biases to our database. To alleviate such potential biases, we derived the species-level trait values by first calculating the median trait-specific value per (sub-)dataset. Then, we calculated the species-level trait averages across those median values. We selected the median instead of mean in this first step as it is less sensitive to outliers and erroneous individual measurements compared to the mean. To further reduce the effect of outlier subdatasets dominating species-level average values, we additionally log-transformed the median values for traits that tend to be highly right-skewed, such as plant height, seed mass, and leaf area. Categorical traits, although most often converging on a single value per species in the source data, was averaged by selecting the most commonly occurring value (i.e., the mode) per species across all datasets.

### 2.5 Phylogenetic distances

To enhance imputation accuracy, we incorporated phylogenetic eigenvectors derived from phylogenetic trees to account for species evolutionary relatedness (Debastiani et al. 2021). Previous comparisons of imputation methodologies have demonstrated that including phylogenetic information significantly improves the estimation of missing trait values and preserves important allometric relationships between traits even when up to 60% of data are missing (Penone et al. 2014, Schrodt et al. 2015).

We used the *V.PhyloMaker2* R package (Jin 2025) to create a phylogenetic tree that includes all our 3099 species. The tree was constructed using the *phylo.maker* function and the default parameter settings (Jin and Qian 2022). The package retrieves a vascular plant megatree where species naming adheres to the Plant List, and we therefore applied the Plant List nomenclature in our species list (see section 2.1) to match the species names with the species in our forming database. The function further prunes the existing tree and attaches any missing species in the megatree based on their higher level taxonomy (i.e., genus and family). Next, we used the pruned tree to calculate phylogenetic eigenvectors using the *PVRdecomp* function from the *PVR* R package (Santos 2018). We selected the 10 first eigenvectors to be used in the imputation model.

### 2.6 Trait Imputation

To estimate likely values for missing traits we conducted trait imputation. A total of 205 trait variables and 10 phylogenetic eigenvectors (columns in the species x traits matrix) were used as input for the imputation process.

We imputed the missing species-level trait values using a Random Forest -based modeling framework, implemented in the *MissForest* R package (Stekhoven and Buhlmann 2012). This non-parametric method is particularly suitable for plant functional trait data due to its ability to simultaneously handle different types of variables (continuous, binary, categorical) and complex interactions between included variables (i.e., the 205 traits and 10 phylogenetic eigenvectors) (Debastiani et al. 2021). In previous studies, the Random Forest approach has shown lower imputation error and bias compared to other imputation methods, especially for data with high proportion of missing values, as was the case for our species-level trait data prior to imputation (Penone et al. 2014, May et al. 2023, Gendre et al. 2024).

The Random Forest -based modelling approach treats the imputation problem as a series of prediction tasks following a systematic and iterative approach, where each variable containing missing values becomes a target variable to be predicted using all other available variables as predictors (Stekhoven and Buhlmann 2012). The *MissForest* algorithm begins by making an initial guess for all missing values in the dataset. Variables are then sorted according to the ascending amount of missing values. For each variable, a random forest model is trained using observed values of the focal variable as a response and all other variables as predictors. The missing values for the focal variable are subsequently predicted using the trained model. This process is repeated iteratively until it halts when a specific criterion is met, namely when the difference between the newly imputed data matrix and the previous one increases for the first time across both continuous and categorical variable types. The performance of the imputation is assessed using the Normalized Root Mean Squared Error (NRMSE) for continuous variables and the Proportion of Falsely Classified entries (PFC) for categorical variables. This method doesn’t require parameter tuning and makes minimal assumptions about data distribution. We set the number of trees to 300 and the minimum size of terminal nodes to 1 for continuous variables and 2 for categorical variables. We used the ‘forest’ mode for parallelization.

Finally, we postprocessed the imputed dataset by backtransforming the log transformed variables that were transformed prior the imputation and by rounding numerical values for traits that originally were integers. At this stage we also removed traits related to biogeography or macroecology and other redundant traits (i.e., when several traits described the same form or functionality, in which case we typically selected the trait with least missing values in the original trait data).

### 2.7 Quality assessment

Our final imputed dataset included 44 traits of which, on average, 56% of species-level trait values were missing before imputation (**Table 3**). Photosynthetic pathway, plant height, and nitrogen fixation capability were the best covered traits while root phosphorus content, specific root area and root dry matter content were the most infrequent. When considering the three most commonly utilized traits, i.e. SLA, plant height, and seed mass (the leaf-height-seed plant ecology strategy scheme; Westoby 1998), 54% of the species had all three traits covered while 21% lacked all the three. However, these numbers shifted to 65% and 6%, respectively, when the megadiverse but complex genera *Ranunculus*, *Hieracium* and *Taraxacum* were excluded.

**Table 2.** Summary of the traits included in the final NordicTraits dataset. In the Data type: B = binomial; C = categorical; N = numerical; O = Ordinal.

| Trait | Explanation | Category | Data type | Unit | Values/Levels | Level description |
| --- | --- | --- | --- | --- | --- | --- |
| dispersal_distance_class | Categorizes how far seeds or propagules are typically dispersed from the parent plant. | dispersal | O | index | min = 1; max = 6 | 1 = dispersal distance for 50 and 99% of seeds 0.1–1 m, 2 = 1–5 m, 3 = 2–15 m, 4 = 40–150 m, 5 = 10–500 m, 6 = 400–1500 m, 7 = 500–5000 |
| dispersal_vector | Describes the primary agent responsible for seed dispersal. | dispersal | C |  | Adh; AirD; AirP; AirW; Bird; Expl; Myr; Passive; Veg; Water | Adh = fruits hooked or sticky, facilitating dispersal in the fur or feathers of animals, AirD = fruits very small and light, thereby easily carried by the wind, AirP = fruits plumed by hairs facilitating wind dispersal, AirW = fruits with wings facilitating wind dispersal, Bird = fruits with a nutritious colored tissue attractive as food of birds (or rarely mammals), Expl = fruits with active explosive/ballistic dispersal mechanisms, Myr = fruits with an elaiosome attracting ants that carry away the fruits, Passive = fruits without functional adaptations to any particular vector, Veg = predominantly vegetative dispersal by stolons, rhizomes, runners, etc., Water = fruits with spongy tissue and enhanced floating ability facilitating water dispersal. |
| seed_buoyancy | Indicates how well seeds float in water, affecting dispersal in aquatic environments. | dispersal | N | index | min = 0.02; max = 1 | 1 indicates that all the seeds float still after a week in still water. 0 means that all seeds sink immediately after placing them in water. |
| seed_terminal_velocity | Measures how fast seeds fall through air, influencing dispersal distance by wind. | dispersal | N | m/s | min = 0.05; max = 5.72 |  |
| palatability | Reflects how attractive or edible the plant is to herbivores. | ecosystem | N | index | min = 1; max = 9 |  |
| growth_form | Describes the overall physical structure or shape of the plant. | growth_form | C |  | Chamaephyte; Cryptophyte; Hemicryptophyte; Liana; Phanerophyte; Therophyte; Vascular parasite; Vascular semi-parasite | Chamaephyte = Perennial plants with buds close to the ground, protected by snow or litter during unfavorable seasons; Cryptophyte = Perennial plants with buds buried underground or underwater, surviving harsh conditions below the surface; Hemicryptophyte = Perennial plants with buds at or just below the soil surface, allowing regrowth after seasonal dieback; Liana = Long-stemmed, woody vines that climb trees or other supports to reach sunlight; Phanerophyte = Woody plants with buds exposed above ground, typically trees or shrubs; Therophyte = Annual plants that complete their life cycle in one season, surviving as seeds during unfavorable periods; Vascular parasite = Plants that attach to and derive nutrients from the vascular system of host plants, often harming them; Vascular semi-parasite = Plants that photosynthesize but also extract water and nutrients from host plants, usually without killing them. |
| growth_form_aquatic | Specifies if the plant grows submerged, | growth_form | O |  | aquatic; semiaquatic; |  |
|  | floating, or emergent in aquatic habitats. |  |  |  | terrestrial |  |
| woodiness | Indicates whether the plant has woody stems or is herbaceous. | growth_form | O |  | non-woody; semi-woody; woody | non-woody = herbaceous plant, semi-woody = herbaceous plant with a woody base of the stem, woody = woody plant |
| LDMC | Leaf Dry Matter Content: the ratio of dry mass to fresh mass in leaves. | leaf_economic | N | g/g | min = 0.045; max = 0.678 |  |
| SLA | Specific Leaf Area: the area of leaf per unit dry mass. | leaf_economic | N | mm<sup>2</sup> mg<sup>-1</sup> | min = 1.15; max = 185.79 |  |
| leaf_C | The concentration of carbon in leaf tissue. | leaf_economic | N | mg/g | min = 144.9; max = 580.1 |  |
| leaf_CN_ratio | The ratio of carbon to nitrogen in leaves, indicating nutrient use efficiency. | leaf_economic | N | ratio | min = 2; max = 100.6 |  |
| leaf_N | The concentration of nitrogen in leaf tissue, important for protein synthesis and growth. | leaf_economic | N | mg/g | min = 2.6; max = 68 |  |
| leaf_NP_ratio | The ratio of nitrogen to phosphorus in leaves, reflecting nutrient limitation. | leaf_economic | N | ratio | min = 3.1; max = 60 |  |
| leaf_P | The concentration of phosphorus in leaf tissue, critical for energy transfer and growth. | leaf_economic | N | mg/g | min = 0; max = 7.7 |  |
| leaf_chlorophyll | The concentration of chlorophyll in leaves, related to photosynthetic efficiency. | leaf_economic | N | mg/g | min = 0.87; max = 28.04 |  |
| longevity | The typical lifespan of the plant. | longevity | O | index | min = 1; max = 4 | 1 = strictly annual, 2 = biennial or at least dying after flowering (hapaxanth), 3 = short lived perennial (often hapaxanthic but may sometimes flower for additional years), 4 = long-lived perennial. |
| fruit_type | Describes the structure and type of fruit produced. | reproduction | C |  | achene; berry; capsule; drupe; follicle; nut; other; pod; siliqua |  |
| nectar_production | Indicates whether and how much nectar the plant produces to attract pollinators. | reproduction | O | index | min = 1; max = 7 | 1 = no nectar production (0 g sugar/m<sup>2</sup>/year) and no collectable pollen, 2 = nectar production insignificant (<0.2 g), or absent but with low but significant amounts of collectable pollen, 3 = nectar production small (0.2–5 g), or lower but with copious collectable pollen 4 = nectar production modest (5–20 g), 5 = rather large (20–50 g), 6 = large (50–200 g), 7 = very large (>200 g). |
| pollinator_type | Specifies the primary type of pollinator the plant attracts. | reproduction | C |  | 0a; 0b; 0c; 0d; 1; 1a; 1ab; 1b; 2; 2a; 2ab; 2b | The data are given on a three-degree scale referring to the degree of dependence on pollinating animals, with subscript letters denoting the kind(s) of pollinating agents concerned: 0a = independent of pollinators, mostly or only vegetatively propagated by fragmentation or clonal growth, 0b = independent of pollinators, mostly or only pollinated by abiotic agents (wind or water; this also includes ferns and fern-allies which are not pollinated but dependent on water for fertilization), 0c = independent of pollinators, mostly or only selfing, 0d = independent of pollinators, mostly or only producing seeds by apomixis (excluding pseudogams that need pollination of the endosperm), 1 = pollination facilitated by insects, in addition to selfing (i.e. self-compatible) and/or pollination by abiotic agents, 2 = exclusively pollinated by insects (i.e. self-incompatible): 1/2a = exclusively pollinated by Hymenoptera (bees and bumblebees), 1/2ab = exclusively pollinated by Hymenoptera and Lepidoptera, 1/2b = exclusively pollinated by Lepidoptera (butterflies and moths), 1/2 = mostly pollinated by insects other than Lepidoptera or Hymenoptera (e.g. Diptera or beetles). |
| seed_bank_persistence | Describes how long seeds remain viable in the soil seed bank. | reproduction | O | index | min = 1; max = 4 | The four-degree scale translates as: 1 = transient (up to one or rarely two years), 2 = short-lived (1–5 years), 3 = long-lived (5–25 years), 4 = semipermanent (>25 years). |
| seed_dormancy | Indicates whether seeds require specific conditions to germinate, affecting timing of growth. | reproduction | O | index | min = 1; max = 4 | The four-degree scale, possibly better treated as categories, translates as: 1 = non-dormant, germinates within 10 days, 2 = non-dormant or with non-deep physiological or physical dormancy, germinates 10–30 days after sowing, 3 = with morphological, physical or mechanical dormancy, or with intermediate physiological dormancy resulting in very slow germination but independent of temperature and season (or requiring <35 days of cold stratification), 4 = with intermediate or deep physiological or morphophysiological dormancy, only germinating after at least one period of > 35 days of cold stratification. |
| seed_longevity_index | Measures how long seeds remain viable under storage conditions. | reproduction | N | index | min = 0; max = 1 |  |
| seed_mass | The mass of a seed, influencing dispersal, establishment, and survival. | reproduction | N | mg | min = 0.37; max = 5268.49 |  |
| RDMC | Root Dry Matter Content: the ratio of dry mass to fresh mass in roots. | root | N | g/g | min = 0.053; max = 0.505 |  |
| SRA | Specific Root Area: the surface area of roots per unit dry mass, linked to nutrient uptake. | root | N | cm<sup>2</sup>/g | min = 0.04; max = 4857.71 |  |
| SRL | Specific Root Length: the length of roots per unit dry mass, related to resource acquisition. | root | N | m/g | min = 0.7; max = 12047.28 |  |
| mycorrhiza_colonization_intensity | The extent to which roots are colonized by mycorrhizal fungi, aiding nutrient uptake. | root | N | % | min = 0; max = 100 |  |
| mycorrhiza_type | Specifies the type of mycorrhizal association. | root | C |  | AM; AM/EM; EM; EriM; EriM/ArbM; MonM; None; OrkM; PyrM | None = no mycorrhizal associations, AM = arbuscular mycorrhiza, AM/EM = with both arbuscular mycorrhiza and ectomycorrhiza, ArbM = arbutoid mycorrhiza, EM = ectomycorrhiza, ErM = ericoid mycorrhiza, MonM = |
|  |  |  |  |  |  | monotropoid mycorrhiza, OrkM = Orchid mycorrhiza, PyrM = Pyrola mycorrhiza. |
| root_N | The concentration of nitrogen in root tissue, important for growth and metabolism. | root | N | mg/g | min = 1.4; max = 40 |  |
| root_P | The concentration of phosphorus in root tissue, critical for energy transfer and growth. | root | N | mg/g | min = 0.45; max = 7.02 |  |
| root_mass_fraction | The proportion of total plant biomass allocated to roots, reflecting belowground investment. | root | N | ratio | min = 0.0039; max = 0.925 |  |
| height | The maximum height the plant typically reaches, influencing light competition and dispersal. | size | N | cm | min = 0.5; max = 1712.1 |  |
| leaf_area | The total surface area of a leaf, related to light interception and gas exchange. | size | N | mm<sup>2</sup> | min = 0.6; max = 62774.2 |  |
| leaf_length | The length of a leaf, influencing light capture and transpiration. | size | N | mm | min = 0.6; max = 423.4 |  |
| leaf_mass | The average mass of a leaf, related to resource investment and function. | size | N | mg | min = 0.4; max = 4055.4 |  |
| leaf_thickness | The thickness of a leaf, affecting light absorption, gas exchange, and drought tolerance. | size | N | mm | min = 0.386; max = 4.047 |  |
| rooting_depth | The maximum depth roots penetrate into the soil, affecting access to water and nutrients. | size | N | m | min = 0.37; max = 5.15 |  |
| Grime_strategy | Classifies plants into competitive, stress-tolerant, or ruderal strategies based on habitat. | strategy | C |  | C; CR; CS; CSR; R; S; SR | The Grime CRS plant strategy system.<br>C (Competitor) = Plants that thrive in productive, low-stress, low-disturbance environments by outcompeting neighbors for resources.<br>R (Ruderal) = Plants adapted to high-disturbance, low-stress environments, characterized by rapid growth and reproduction.<br>S (Stress-tolerator) = Plants adapted to low-resource, high-stress environments, growing slowly but persisting under harsh conditions.<br>Some species may exhibit mixed strategies (e.g., CS, CR, or CSR), reflecting adaptations to intermediate conditions. |
| carnivory | Indicates whether the plant obtains nutrients by trapping and digesting animals. | strategy | B |  | no; yes | The ability (yes), or inability (no) to trap and digest animals. |
| leaf_phenology | Describes the seasonal timing of leaf production, growth, and senescence. | strategy | O |  | deciduous; variable; evergreen | deciduous = plants shed all their leaves seasonally, usually in response to cold or dry periods; variable = plants that can exhibit either deciduous or evergreen traits, depending on environmental conditions or species variability; evergreen = plants retain their leaves year-round, replacing them gradually. |
| nitrogen_fixation | Indicates whether the plant can convert atmospheric nitrogen into a usable form via symbiosis. | strategy | B |  | no; yes | The ability (yes), or inability (no) to form symbioses with bacteria capable of fixing atmospheric nitrogen |
| parasitism | Describes whether the plant obtains resources by parasitizing other plants. | strategy | O |  | none; hemiparasite; holoparasite | none = non-parasitic species, hemiparasite = own photosynthesis but extracting water and mineral nutrients from the host, holoparasite = fully dependent on the host also for carbohydrates. |
| photosynthetic_pathway | Specifies the photosynthetic pathway used, affecting water and carbon use. | strategy | C |  | C3; C4; CAM | C3 = the most common pathway, where CO<sub>2</sub> is fixed directly into a 3-carbon compound. Efficient in cool, moist conditions but prone to photorespiration in hot, dry climates; C4 = CO<sub>2</sub> is first fixed into a 4-carbon compound, reducing photorespiration. Common in hot, sunny environments; CAM |
|  |  |  |  |  |  | (Crassulacean Acid Metabolism) = CO<sub>2</sub> is fixed at night and stored as an acid, then used during the day to minimize water loss. Typical in arid-adapted plants. |

**Table 3.** Statistics of the quality assessment. Proportion of species covered indicates the share of species that were present in the original data for a given trait. Phylogenetic signal (Lambda) was calculated both for the species-level trait values (raw) and the missingness of the values across species. A high Lambda value indicates highly aggregated values in phylogenetic space. R = Spearman correlation coefficient; NRMSE = Normalized root mean squared error; PCC = Proportion of correctly classified. R, NRMSE, PCC and Kappa represent the evaluation statistics from the cross validation procedures. The statistics without the “sp” ending relate to cross validation where a random sample of 1% of the species-level trait values were excluded from the imputation rounds. The statistics with the “sp” ending relate to cross validation where a random set of 1% of the species were excluded from the trained imputation model in each iteration.

| Trait | Proportion of species covered | Phylogenetic signal raw | Phylogenetic signal missingness | R | NRMSE | PCC | Kappa | R_sp | NRMSE_sp | PCC_sp | Kappa_sp |
| --- | --- | --- | --- | --- | --- | --- | --- | --- | --- | --- | --- |
| dispersal_distance_class | 0.67 | 0.97 | 0.79 | 0.95 | 0.36 |  |  | 0.38 | 1.07 |  |  |
| dispersal_vector | 0.73 | 0.97 | 0.84 |  |  | 0.90 | 0.86 |  |  | 0.61 | 0.31 |
| seed_buoyancy | 0.24 | 0.74 | 0.47 | 0.52 | 0.86 |  |  | 0.48 | 0.88 |  |  |
| seed_terminal_velocity | 0.30 | 0.96 | 0.55 | 0.84 | 0.57 |  |  | 0.35 | 0.95 |  |  |
| palatability | 0.19 | 0.88 | 0.46 | 0.74 | 0.71 |  |  | 0.69 | 0.80 |  |  |
| growth_form | 0.47 | 0.92 | 0.66 |  |  | 0.86 | 0.78 |  |  | 0.58 | 0.16 |
| growth_form_aquatic | 0.71 | 0.87 | 0.84 |  |  | 0.98 | 0.74 |  |  | 0.96 | 0.00 |
| woodiness | 0.57 | 0.95 | 0.71 |  |  | 0.97 | 0.84 |  |  | 0.90 | 0.04 |
| LDMC | 0.54 | 0.90 | 0.72 | 0.84 | 0.56 |  |  | 0.51 | 0.87 |  |  |
| SLA | 0.60 | 0.81 | 0.76 | 0.74 | 0.67 |  |  | 0.24 | 1.07 |  |  |
| leaf_C | 0.25 | 0.68 | 0.51 | 0.61 | 0.81 |  |  | 0.14 | 1.25 |  |  |
| leaf_CN_ratio | 0.22 | 0.54 | 0.57 | 0.68 | 0.76 |  |  | 0.39 | 0.96 |  |  |
| leaf_N | 0.34 | 0.76 | 0.60 | 0.74 | 0.68 |  |  | 0.46 | 0.91 |  |  |
| leaf_NP_ratio | 0.18 | 0.74 | 0.63 | 0.46 | 0.90 |  |  | 0.11 | 1.02 |  |  |
| leaf_P | 0.26 | 0.66 | 0.60 | 0.66 | 0.77 |  |  | 0.22 | 1.05 |  |  |
| leaf_chlorophyll | 0.11 | 0.69 | 0.49 | 0.54 | 0.84 |  |  | 0.25 | 0.97 |  |  |
| longevity | 0.73 | 0.88 | 0.81 | 0.93 | 0.38 |  |  | 0.23 | 1.06 |  |  |
| fruit_type | 0.37 | 0.82 | 0.81 |  |  | 0.89 | 0.83 |  |  | 0.73 | 0.55 |
| nectar_production | 0.71 | 0.99 | 0.88 | 0.91 | 0.42 |  |  | 0.68 | 0.76 |  |  |
| pollinator_type | 0.67 | 0.83 | 0.85 |  |  | 0.71 | 0.64 |  |  | 0.49 | 0.37 |
| seed_bank_persistence | 0.44 | 0.84 | 0.62 | 0.74 | 0.69 |  |  | 0.17 | 1.04 |  |  |
| seed_dormancy | 0.34 | 0.84 | 0.74 | 0.71 | 0.71 |  |  | 0.35 | 1.04 |  |  |
| seed_longevity_index | 0.38 | 0.60 | 0.59 | 0.65 | 0.78 |  |  | 0.15 | 1.05 |  |  |
| seed_mass | 0.65 | 0.99 | 0.81 | 0.86 | 0.53 |  |  | 0.31 | 0.97 |  |  |
| RDMC | 0.07 | 0.74 | 0.52 | 0.40 | 0.91 |  |  | -0.03 | 1.09 |  |  |
| SRA | 0.06 | 0.23 | 0.38 | 0.67 | 0.78 |  |  | -0.13 | 1.05 |  |  |
| SRL | 0.14 | 0.11 | 0.50 | 0.39 | 0.92 |  |  | 0.02 | 1.06 |  |  |
| mycorrhiza_colonization_intensity | 0.38 | 0.88 | 0.57 | 0.77 | 0.65 |  |  | 0.64 | 0.78 |  |  |
| mycorrhiza_type | 0.72 | 0.98 | 0.86 |  |  | 0.96 | 0.92 |  |  | 0.87 | 0.72 |
| root_N | 0.13 | 0.73 | 0.54 | 0.63 | 0.82 |  |  | 0.45 | 0.89 |  |  |
| root_P | 0.05 | 0.78 | 0.48 | 0.44 | 0.90 |  |  | -0.03 | 1.00 |  |  |
| root_mass_fraction | 0.14 | 0.68 | 0.54 | 0.58 | 0.82 |  |  | 0.32 | 1.00 |  |  |
| height | 0.78 | 0.94 | 0.84 | 0.92 | 0.41 |  |  | 0.19 | 1.13 |  |  |
| leaf_area | 0.06 | 0.95 | 0.78 | 0.94 | 0.37 |  |  | 0.42 | 0.97 |  |  |
| leaf_length | 0.35 | 0.86 | 0.75 | 0.84 | 0.56 |  |  | 0.41 | 0.97 |  |  |
| leaf_mass | 0.54 | 0.93 | 0.71 | 0.92 | 0.41 |  |  | 0.42 | 0.96 |  |  |
| leaf_thickness | 0.22 | 0.41 | 0.58 | 0.66 | 0.76 |  |  | 0.42 | 0.92 |  |  |
| rooting_depth | 0.20 | 0.28 | 0.62 | 0.66 | 0.78 |  |  | 0.19 | 0.98 |  |  |
| Grime_strategy | 0.55 | 0.66 | 0.79 |  |  | 0.96 | 0.94 |  |  | 0.35 | 0.01 |
| carnivory | 0.75 | 1.00 | 0.84 |  |  | 1.00 | 0.50 |  |  | 1.00 | 0.00 |
| leaf_phenology | 0.05 | 0.51 | 0.72 |  |  | 0.76 | 0.43 |  |  | 0.61 | 0.08 |
| nitrogen_fixation | 0.75 | 0.97 | 0.87 |  |  | 1.00 | 0.98 |  |  | 1.00 | 1.00 |
| parasitism | 0.73 | 1.00 | 0.83 |  |  | 1.00 | 0.90 |  |  | 0.99 | 0.00 |
| photosynthetic_pathway | 0.81 | 0.96 | 0.75 |  |  | 0.98 | 0.35 |  |  | 0.98 | 0.00 |

We calculated the phylogenetic signal of missingness for each trait, i.e., how strongly the missing values were linked to evolutionary relationships across species. The index was calculated using the *phylosig* function from the *phytools* R package (Revell 2024) by applying the lambda method (Pagel 1999) based on the pruned phylogenetic tree described above. A high lambda value indicates that missing values are phylogenetically highly clustered, suggesting that the imputation process has to extrapolate in evolutionary space. Lambda of missingness was highest in nectar production and lowest in specific root area (**Table 3**). We also calculated the phylogenetic signal in the species-level trait values used in the imputation. Here, carnivory and parasitism showed the highest signal and specific root length the lowest. Users of the NordicTraits are advised to exercise caution when using imputed values for traits with both an overall high proportion of missing values and a strong phylogenetic signal of missing values ‒ especially when a strong phylogenetic signal in the trait values themselves was also detected.

To evaluate an imputation output, it is advised to conduct a careful leave-one-out cross validation (Gorné et al. 2025). However, the size of our dataset means that running the MissForest procedure once would take 5-7 hours even when parallelized with 12 computing cores. Thus, the most robust cross validation was not computationally feasible. Instead, we developed two additional cross validation (CV) procedures that require less computational power.

1. To evaluate imputation robustness and output reliability for data-rich species, we assigned part of the data as missing values and imputed their values. For each trait we randomly selected 1% of the species-level trait values and set them to NoData before the imputation, i.e., set their actual observations aside as a testing dataset. We ran the imputation model on the version where we had artificially assigned values as NoData and compared the imputed values to the true values. This process was repeated 20 times with different sets of randomly selected 1% of species-level trait values of each trait.
2. To evaluate imputation robustness and output reliability for data-poor species for which only phylogenetic information is available for imputation, we randomly selected 1% of the species and set all their trait values as NoData before the imputation. We ran the imputation model and compared the imputed values to their observed values and repeated this process 20 times.

In both of the above approaches, we calculated the normalized root mean squared error (NRMSE) and Spearman correlation coefficient (R) for continuous traits and used these statistics for evaluation. For categorical and binary trait predictions we used the proportion of correctly classified values and Cohen’s Kappa statistic to evaluate goodness of fit. All evaluation statistics from the cross validation procedures are presented in **Table 3**. The average correlation between the observed and predicted numeric traits was 0.71 for the first type of CV but dropped to 0.30 in the second CV procedure. The average percentage of correctly classified statistics showed a similar pattern for categorical traits, 0.92 and 0.72 for the first and second CV approach, respectively. This clearly indicates that the trait-trait correlations are crucial information for the imputation, and that the phylogenetic Eigenvectors alone provide much poorer trait predictions than when used in combination with trait-trait correlations. In particular, root traits, which were originally underrepresented in the source data, exhibited very low predictive accuracy during imputation when only phylogenetic data were available for a given species, suggesting that imputation results of root traits were unreliable for species with the fewest observed traits and should be used with caution.

## 3. Format of the dataset

The NordicTraits dataset consists of six data table files that can be accessed in the Supplementary materials of this article. The six files are as follows:

1. *NordicTraits_wide_V1.csv:* An imputed dataset of species-level trait values in a wide format as a comma-limited csv file. This is a gap-free species x traits matrix where rows represent the 3099 species and columns the 44 traits.
2. *NordicTraits_long_V1.csv:* An imputed dataset of species-level trait values in a long format as a comma-limited csv file. Here, the same imputed trait information is presented in the long format where all trait names are under column “*trait*” and trait values under “*values*”. Additionally, users can see which species-level trait values were imputed and which were not (column “*imputed*”) as well as their units and the main trait category that they belong to.
3. *NordicTraits_metadata_traits_V1.xlsx:* Metadata explaining the traits as an xlsx file (extended version of the **Table 2**).
4. *NordicTraits_metadata_species_V1.xlsx:* Species metadata as an xlsx file (table that matches our final nomenclature to the species names used in the national species lists as well as the national taxon ID numbers and vernacular names when available).
5. *NordicTraits_evaluation_statistics_V1.xlsx:* Validation statistics as an xlsx file (extended version of the **Table 3**).
6. *NordicTraits_metadata_traits_extended_V1.xlsx:* Metadata showing all the 205 traits included in the imputation model in an xlsx file.

The NordicTraits dataset is planned to be updated at irregular intervals. The most recent version can always be checked and accessed in the project’s GitHub repository which also includes the computer code used to produce the dataset: https://github.com/poniitty/NordicTraits

## 4. Discussion

The NordicTraits dataset provides the first comprehensive, gap-free, and openly-available species-level functional trait resource for all native vascular plants across Denmark, Finland, Iceland, Norway, and Sweden. We compiled millions of trait measurements and hundreds of traits, performed careful taxonomic harmonization, and rigorous data preprocessing and filtering. Moreover, we used a robust Random Forest based imputation method taking into account phylogenetic relationships to predict missing trait values. As a result, NordicTraits covers a highly diverse set of 44 functional traits for 3099 species. The dataset addresses critical gaps in the availability of trait data that until now have limited trait-based analyses in the Nordic region. Many trait-based statistical approaches require complete datasets with no missing values across the species and traits investigated, which makes this imputed dataset particularly valuable. More fundamentally, our dataset greatly expands the current understanding of the plant trait space and its axes of variation for the entire Nordic species pool (Diaz et al. 2016), with potential for breakthrough discoveries e.g. in the relationships between plant function and the environment (Joswig et al. 2022) or the threats that plant functional diversity is facing under anthropogenic pressures (Carmona et al. 2021).

It is important to note, however, that NordicTraits is a dataset based on modelling. Even the most advanced imputation techniques, such as those used here, inherit the sparsity of global trait matrices. The imputation approach alleviates practical issues caused by gaps in the original data but it cannot recover information where it is completely missing. This means that care should be paid when utilizing the imputed values for less well-studied species, taxonomic groups, or regions (Swenson 2014, Hortal et al. 2015b, Grenié et al. 2025). The reliability of these imputed values is still dependent on the amount and pattern of missing values and the correlative structures of the available data (Gorné et al. 2025). NordicTraits has not been curated in detail, and therefore, the trait values should be used with caution. This dataset will likely be most useful in applications where a large number of species and traits are considered and the absolute accuracy of a single value is less crucial (Gorné et al. 2025). In contrast, studies that consider only a few species or species that are ecologically similar, or studies where the focus is on specific traits, should carefully consider the limitations of this dataset before using it.

NordicTraits contains species-level trait data. This means that it does not consider intraspecific trait variation and also lacks information on the geographic origin of single observations of trait values. Earlier studies have shown that plant height, leaf nutrient contents, and root structural traits in particular exhibit substantial intraspecific variation (Siefert et al. 2015, Firn et al. 2019, Rodríguez-Alarcón et al. 2024). NordicTraits is based on vast amounts of source data for a wide range of species, involving a trade-off between coverage and generalizability on the one hand, and detail and geographic explicitness on the other hand. As species with broad environmental ranges often exhibit significant intraspecific variation, depending on the geographic origin of the original measurements, the raw data may disproportionately represent, e.g., the warmer or colder extremes of their distribution. Such geographic bias can skew mean trait values and thereby affect related species rankings in the imputation process. Therefore, users focusing on specific regions, such as the Arctic tundra, should critically assess whether the trait values in this dataset accurately reflect their focal populations. For example, traits for a true Arctic species are likely sourced from that ecosystem, while species with large geographic ranges may be represented by data from warmer areas, potentially overestimating traits like size or height in marginal Arctic populations (Kemppinen and Niittynen 2022). Also, it is important to acknowledge that, although this dataset covers plant species occurring in the Nordic countries, the trait observations may often stem from populations outside of this region. Thus, we recommend careful evaluation when applying these data to localized studies.

The trait values that NordicTraits provides are imputed values. Imputed trait values are not the same as real trait measurements (Gorné et al. 2025), and this should be kept in mind when using NordicTraits. If imputed values are treated as real measurements in subsequent analyses, this may produce standard errors, confidence intervals, and p-values that are unrealistically small, compared to an analysis that is based only on observed data (i.e., the single imputation fallacy (Blomberg and Todorov 2025)). Using multiple imputations can more robustly account for uncertainty in the imputed values, but would be impractical to include in a published dataset like NordicTraits due to the high number of species it contains. Thus, we encourage the user to employ the provided estimates of uncertainty in the output and carefully consider their implications for the user’s applications.

While compiling NordicTraits, we experienced first-hand the ubiquitous gaps in openly available trait data. Firstly, the documentation and metadata provided in the original sources was often limited, insufficient, or even non-existing. Thus, it was difficult to evaluate how the methodological choices may have influenced the trait measurements. Secondly, some trait categories, such as below-ground traits or clonal traits, are clearly underrepresented in trait literature and databases. These underrepresented traits cover only a small proportion of the species even in the Nordic countries, which likely have better overall availability of trait data compared to many other regions in the World (Grenié et al. 2025). Thirdly, the main body of the trait measurements cover the most common and abundant species. This may not pose a problem in studies on, e.g., ecosystem functioning, but can hinder the applicability of trait-based approaches in conservation and biodiversity research. Thus, we recommend that future research involving collection on trait measurements would 1) put effort in assuring high quality of metadata, 2) gather information also on less commonly measured traits, and 3) prioritize targeted data collection to expand taxonomic (and geographical) coverage. Overall, we call on trait-based research to make use of standard protocols designed for trait data collection (Cornelissen et al. 2003, Pérez-Harguindeguy et al. 2013).

## 5. Conclusions

The NordicTraits dataset is an important step forward in trait-based ecological research regarding the northern European flora, offering an openly-available and unprecedented resource for understanding plant functional strategies and diversity across the Nordic region. This dataset offers a compiled, harmonised, and imputed set of trait values for 3,099 vascular plant species. The dataset addresses a long-standing gap in regional trait availability and enables researchers to explore mechanistic links between plant strategies, environmental gradients, and ecosystem processes. The broad taxonomic and trait coverage of this dataset will support applications ranging from biodiversity conservation and invasive species management to predictive modeling of vegetation responses to climate and land-use change. The dataset is best suited for macroecological analyses, comparative studies, and broad-scale modeling, where the strengths of a comprehensive, standardized trait matrix outweighs the uncertainties inherent in aggregated and imputed data. As the field of trait-based ecology evolves, datasets like NordicTraits will advance ecological and evolutionary understanding in a rapidly warming Nordic region by supporting broader and more mechanistic research on biodiversity and ecosystem functioning.

## Data Availability Statement

The NordicTraits dataset consists of six data table files that can be accessed in the Supplementary materials of this article. The up-to-date version will be maintained and can be accessed in the project’s GitHub repository which also includes the computer code used to produce the dataset: https://github.com/poniitty/NordicTraits

## Acknowledgements

PN acknowledges funding from the Research Council of Finland (grant no. 378397; 347558; PROFI8: 365202). JK acknowledges funding from the Research Council of Finland (349606; 353218; 370245). MHH acknowledges funding from the Research Council of Finland (grant no. 360742). RKH acknowledges the support by NordForsk through the funding to Adapting LAw for MOving Targets: Climate Change, Overtourism and Biodiversity in Indigenous Arctic National Parks (project number 216938). MHH, A-MM and RKH also acknowledge funding by the Strategic Research Council of Finland through the JUST TRANSITION program for the project MUST (grant 358367). VN acknowledges funding from the Research Council of Finland (grant no. 274489; 345733). A-MM acknowledges funding from the Kvantum Institute at the University of Oulu. We thank all the researchers who have collected and openly provided the original trait data utilized in preparation of the NordicTraits dataset.

